## Supplemental materials for "Network potential identifies therapeutic *miRNA* cocktails in Ewing sarcoma"

### 3.1 Additional Analyses

First, we provide some additional summary plots describing mRNA expression and network potential in our samples.

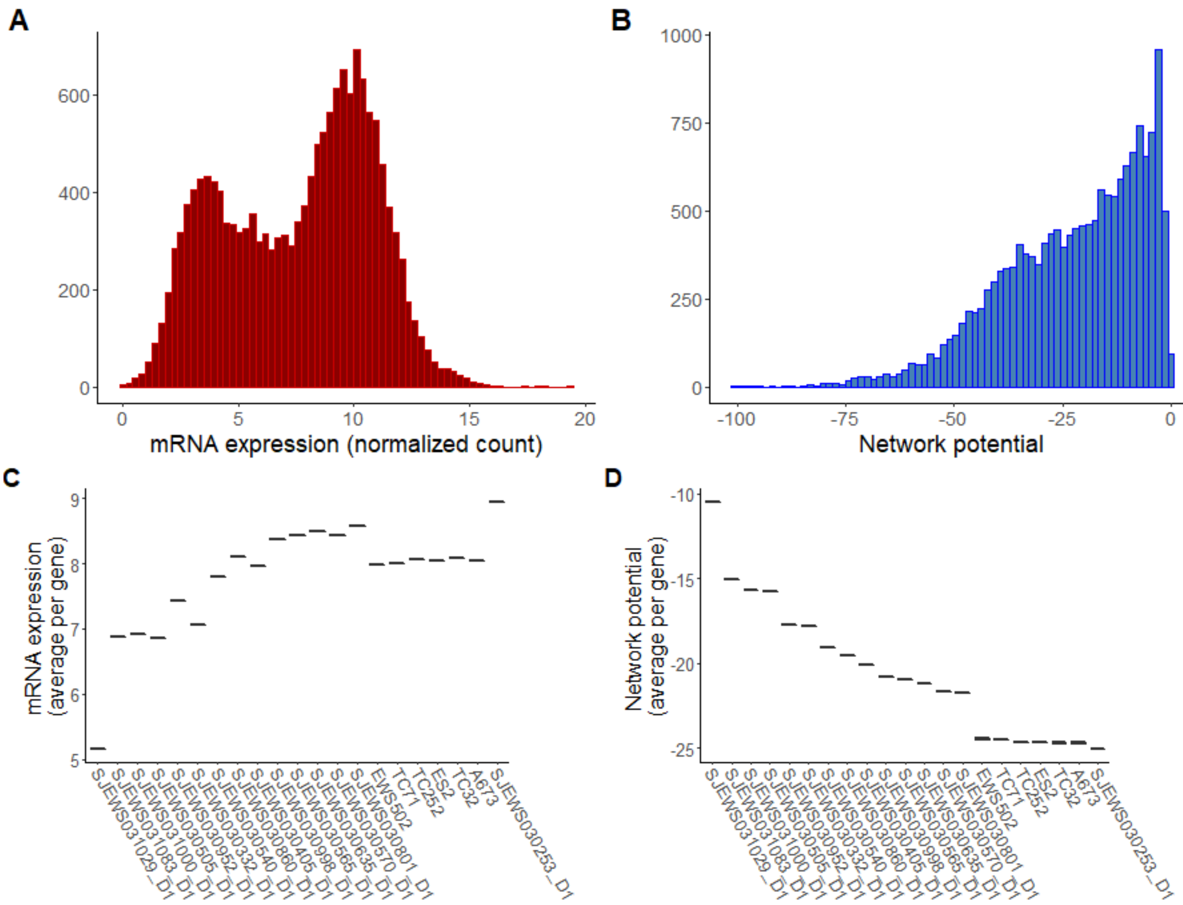


**Figure S1. Network potential describes different features of a cell signaling network compared to mRNA expression alone. Panel A:** Histogram of mRNA expression for each gene (averaged across all samples). **Panel B:** Histogram of the network potential for each gene (averaged across all samples) mRNA transcripts with an expression level of zero were excluded from both histograms to better visualize the distribution of genes that are expressed. **Panel C:** Box plot showing the total mRNA expression for each cell line and patient sample (patient samples begin with SJEWS). **Panel D:** Box plot showing the total network potential for each cell line and patient sample.

As described in the main text, we ranked proteins according to their contribution to network stability by calculating the change in network potential following complete *in silico* repression of each protein. In the main text, we limited our analysis to proteins that had been causally implicated in cancer according to the cosmic database[41]. Here, we present the top 50 proteins (when network potential for all 6 cell lines was averaged) ranked by contribution to network stability, not limited to proteins that were causally implicated in cancer (Figure 4).

We also analyzed each cell line individually to identify the top protein targets for each cell line. In the main text, we limited this analysis to proteins that had been causally implicated in cancer[41]. Here, we present the top protein targets for each cell line, not limited to those proteins that had previously been causally implicated in cancer (Table S1).

To evaluate the impact of PPI choice on our findings, we repeated the protein target selection portion of our analysis using an entirely different PPI, stringdb (**Figure S3**). To ensure a similar number of total edges to the biogrid database, we modulated the provided "interaction score" until the resulting stringdb derived network contained about 400000 edges. We ultimately used an interaction score cutoff of 700 (indicating a high confidence in the observed interaction).

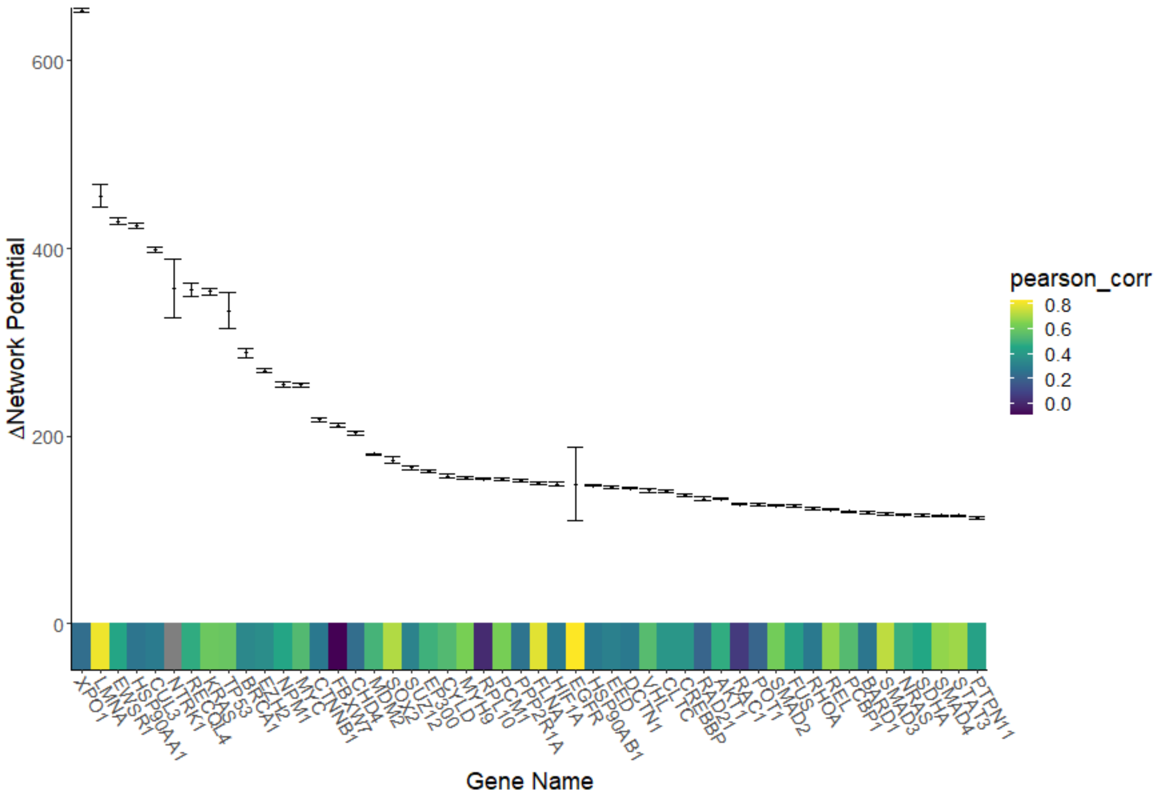


**Figure S2. Protein targets ranked by contribution to network stability.** When averaging across cell lines, XPO1, LMNA, EWSR1, HSP90AA1, and CUL3 were identified as the most important proteins in the Ewing sarcoma cell signaling network (when limiting our analysis to proteins causally implicated in cancer[41]). When each protein was simulated as completely repressed *in silico*, network potential was increased by 654, 456, 429, 425, and 399, respectively. The heatmap at the bottom of the plot describes the protein-mRNA correlation for each gene in the cancer cell line atlas. Grey indicates no data was available. It is reassuring that EWSR1, the kinase associated with Ewing sarcoma development, is identified as highly influential in the cell signaling network by this method.

**Gene set enrichment analysis** We conducted gene set enrichment analysis, using all genes in our Ewing sarcoma cell signaling network as the gene set. We ranked this set of genes by change in network potential and used the "hallmarks" pathways set from the Molecular Signatures Database as the genomic background[43,44]. We also included a gene set corresponding to the miRNA biogenesis pathway[59]. We used the "fgsea" R package version 1.8.0 to conduct the analysis using the following settings: nperm = 500, minSize = 1, maxSize = $\infty$, nproc = 0, gseaParam = 1, BPPARAM =NULL[45]. We found our gene set to be significantly enriched in several pathways related to oncogenesis, including DNA repair, apoptosis, and MTOR signaling (Table S3). These results indicate that network potential can identify a cancer-specific signal from mRNA expression data.

| | TC252 | ES2 | A673 | TC32 | EWS502 | TC71 |
|---|---|---|---|---|---|---|
| 1 | TRIM25 | TRIM25 | TRIM25 | TRIM25 | TRIM25 | TRIM25 |
| 2 | APP | APP | APP | APP | APP | APP |
| 3 | ELAVL1 | ELAVL1 | ELAVL1 | ELAVL1 | ELAVL1 | ELAVL1 |
| 4 | RNF4 | RNF4 | RNF4 | RNF4 | RNF4 | RNF4 |
| 5 | HNRNPL | HNRNPL | HNRNPL | HNRNPL | HNRNPL | HNRNPL |
| 6 | XPO1 | XPO1 | XPO1 | XPO1 | XPO1 | XPO1 |
| 7 | NXF1 | NXF1 | NXF1 | NXF1 | NXF1 | NXF1 |
| 8 | UBC | TNIP2 | UBC | UBC | UBC | UBC |
| 9 | TNIP2 | UBC | TNIP2 | TNIP2 | TNIP2 | TNIP2 |
| 10 | MOV10 | MOV10 | MOV10 | MOV10 | MOV10 | MOV10 |

**Table S1. Top protein targets for each cell line.** We ranked potential targets by predicted change in network potential when each protein was modeled as repressed.

| | TC252 | ES2 | A673 | TC32 | EWS502 | TC71 |
|---|---|---|---|---|---|---|
| 1 | **XPO1** | **XPO1** | **XPO1** | **XPO1** | **XPO1** | **XPO1** |
| 2 | **LMNA** | **LMNA** | **LMNA** | **LMNA** | NTRK1 | EWSR1 |
| 3 | EWSR1 | **HSP90AA1** | **HSP90AA1** | EWSR1 | **HSP90AA1** | LMNA |
| 4 | **HSP90AA1** | **EWSR1** | **EWSR1** | **HSP90AA1** | **EWSR1** | **HSP90AA1** |
| 5 | **CUL3** | **CUL3** | **CUL3** | **CUL3** | LMNA | **CUL3** |
| 6 | **NTRK1** | KRAS | **NTRK1** | **NTRK1** | CUL3 | RECQL4 |
| 7 | **TP53** | **TP53** | RECQL4 | **TP53** | RECQL4 | KRAS |
| 8 | RECQL4 | RECQL4 | KRAS | KRAS | TP53 | NTRK1 |
| 9 | KRAS | EGFR | TP53 | RECQL4 | KRAS | BRCA1 |
| 10 | **BRCA1** | **BRCA1** | **BRCA1** | **BRCA1** | **BRCA1** | TP53 |

**Table S2. Top cancer-associated protein targets for each cell line.** We ranked potential targets by predicted change in network potential when each protein was modeled as repressed, limited to proteins causally associated in cancer according to the Cosmic database. Proteins that appear in the same position for $\geq 3$ cell lines are **bolded**.

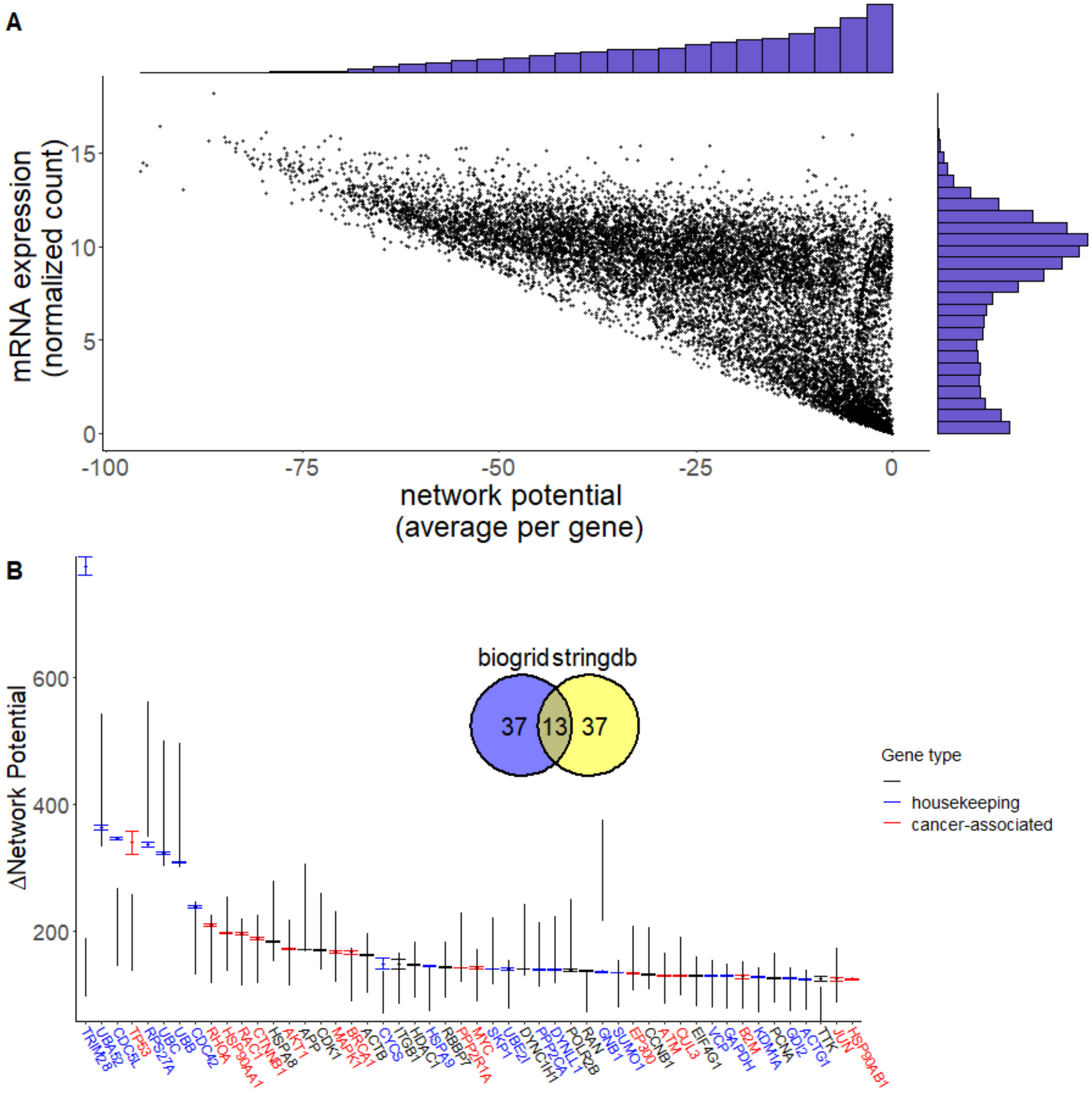


**Figure S3.** **Overview of our findings, using the stringdb protein-protein interaction network. Panel A:** Scatterplot with marginal histograms comparing mRNA expression to network potential. **Panel B:** Box and whisker plot showing the change in network potential for the top 50 genes, as well as 99.99% confidence intervals from the permutation test. We also show a histogram comparing the top 50 genes identified by our pipeline using stringdb compared to biogrid as the protein-protein interaction network.

| | pathway | pval | padj | ES | NES | nMoreExtreme | size |
|---|---|---|---|---|---|---|---|
| 1 | MITOTIC_SPINDLE | 0.00 | 0.01 | 0.62 | 1.47 | 0.00 | 197 |
| 2 | DNA_REPAIR | 0.00 | 0.01 | 0.59 | 1.37 | 0.00 | 146 |
| 3 | G2M_CHECKPOINT | 0.00 | 0.01 | 0.72 | 1.68 | 0.00 | 187 |
| 4 | APOPTOSIS | 0.00 | 0.01 | 0.62 | 1.44 | 0.00 | 158 |
| 5 | PROTEIN_SECRETION | 0.00 | 0.01 | 0.63 | 1.44 | 0.00 | 94 |
| 6 | APICAL_SURFACE | 0.00 | 0.01 | 0.73 | 1.57 | 0.00 | 42 |
| 7 | UNFOLDED_PROTEIN_RESPONSE | 0.00 | 0.01 | 0.62 | 1.43 | 0.00 | 106 |
| 8 | PI3K_AKT_MTOR_SIGNALING | 0.00 | 0.01 | 0.69 | 1.58 | 0.00 | 104 |
| 9 | MTORC1_SIGNALING | 0.00 | 0.01 | 0.61 | 1.43 | 0.00 | 193 |
| 10 | E2F_TARGETS | 0.00 | 0.01 | 0.72 | 1.69 | 0.00 | 195 |
| 11 | MYC_TARGETS_V1 | 0.00 | 0.01 | 0.80 | 1.89 | 0.00 | 193 |
| 12 | OXIDATIVE_PHOSPHORYLATION | 0.00 | 0.01 | 0.61 | 1.42 | 0.00 | 184 |
| 13 | ALLOGRAFT_REJECTION | 0.00 | 0.01 | 0.54 | 1.27 | 0.00 | 191 |
| 14 | MIRNA_BIOGENESIS | 0.00 | 0.01 | 0.80 | 1.73 | 0.00 | 40 |
| 15 | WNT_BETA_CATENIN_SIGNALING | 0.01 | 0.02 | 0.68 | 1.47 | 2.00 | 42 |
| 16 | ANGIOGENESIS | 0.01 | 0.02 | 0.72 | 1.53 | 2.00 | 34 |
| 17 | TGF_BETA_SIGNALING | 0.01 | 0.03 | 0.65 | 1.43 | 4.00 | 53 |
| 18 | MYC_TARGETS_V2 | 0.01 | 0.03 | 0.64 | 1.41 | 4.00 | 58 |
| 19 | P53_PATHWAY | 0.01 | 0.03 | 0.51 | 1.19 | 5.00 | 194 |
| 20 | UV_RESPONSE_UP | 0.01 | 0.04 | 0.53 | 1.23 | 6.00 | 153 |
| 21 | SPERMATOGENESIS | 0.02 | 0.04 | 0.54 | 1.24 | 7.00 | 126 |
| 22 | ADIPOGENESIS | 0.02 | 0.04 | 0.50 | 1.18 | 8.00 | 191 |
| 23 | INTERFERON_GAMMA_RESPONSE | 0.02 | 0.04 | 0.50 | 1.18 | 8.00 | 193 |
| 24 | INTERFERON_ALPHA_RESPONSE | 0.02 | 0.05 | 0.56 | 1.28 | 10.00 | 91 |

**Table S3. Genes ranked by network potential are enriched for several biological pathways related to cancer as well as the miRNA bio-genesis pathway** Pathways with an adjusted p-value $< 0.05$ are shown above. “ES” refers to enrichment score and “NES” refers to the normalized enrichment score. “nMoreExtreme” refers to the number of random gene sets (out of 500) that were more enriched than the test set. Size refers to the number of genes in the pathway that were also present in our mRNA expression dataset.
